# BioTEA: containerized methods of analysis for microarray-based transcriptomics data

**DOI:** 10.1101/2022.05.26.493502

**Authors:** Luca Visentin, Luca Munaron, Federico Alessandro Ruffinatti

## Abstract

Tens of thousands of gene expression data sets describing a variety of model organisms in a many different pathophysiological conditions are currently stored in publicly available databases such as Gene Expression Omnibus (GEO) and ArrayExpress. As microarray technology is giving way to RNA-Seq, it becomes strategic to develop high-level tools of analysis to preserve access to this huge amount of information through the most sophisticated methods of data preparation and processing developed over the years, while ensuring at the same time the reproducibility of the results. To meet this need, here we present bioTEA, a novel software tool that combines the ease of use with the versatility and power of an R/Bioconductor-based differential expression analysis, starting from raw data retrieval and preparation to gene annotation. BioTEA is an R-coded pipeline, wrapped in a Python-based command line interface and containerized with Docker technology. The user can choose among multiple options—including gene filtering, batch effect handling, sample pairing, statistical test type—to adapt the algorithm flow to the structure of the particular data set. All these options are saved in single text file which can be easily shared between different laboratories to deterministically reproduce the results. In addition, a detailed log file provides accurate information about each step of analysis. Overall, these features make bioTEA an invaluable tool for both bioiformaticians and wet-lab biologists interested in transcriptomics.

## 1. Introduction

Gene expression DNA microarrays [1] were introduced in mid-1990s and represented the first cost-effective *-omic* technology for the transcriptome profiling [2]. Such a technology consisted in the hybridization of fluorescently labeled cDNA with solid microchips exposing a collection of thousands of known short DNA sequences attached to the chip surface in a defined position and serving as probes. Commercial and custom-designed microarrays evolved through two main generations of chips: the first *two-color* (or *two-channel*) spotted microarrays [3] and the next *single-channel* (or *one-color*) high-density microarrays [4], mainly popularized by Affymetrix (Santa Clara, California) and capable of a much more reliable genome-wide quantification of the expression changes across multiple samples and different experimental conditions. The need for acquiring, processing, and analyzing the ensuing high-throughput transcriptomics data streams has been a powerful drive for the development of innovative computational methods and statistics approaches within the context of the bioinformatics across the postgenomic era [5–9]. The birth and the rapid growth of the Bioconductor project [10], together with the two most important public repositories for gene expression data – namely the Gene Expression Omnibus (GEO) [11,12] and ArrayExpress (AE) [13,14] – is probably what best embodies the deep impact of that first technological revolution in *-omics* disciplines.

However, starting from the late 2000s, the nascent Next Generation Sequencing (NGS) technology fostered a second paradigm shift in transcriptomics, leading to the development of the RNA-Seq technique [15], the current standard for transcriptome quantification. RNA-Seq is increasingly taking hold because by sequencing, aligning, and counting cDNA fragment reads it makes it possible to overcome many of the limitations inherent to microarrays, among which the necessary dependence on known genome sequences, a relatively high background signal (leading in turn to a poor limit of detection), a narrow (2-3 orders of magnitude) dynamic range due to the saturation of the fluorescent spots, and the challenging normalization procedures needed to compare expression levels among different experiments.

As a consequence, starting from 2015, scientific publications referring to expression microarrays are on a downward trend, having been overtaken around the same year by the increasing number of RNA-Seq-related papers, which, to date, are still in a phase of exponential growth [16]. These publishing trends clearly show that, because of its higher accuracy and thanks to the constant lowering of NGS prices, RNA-Seq is likely to completely replace microarray technology sooner rather than later and, as a result, all the related methods of data analysis are meant to change accordingly. In particular, while many high-level algorithms developed to analyze microarray data (i.e., starting from the normalized expression matrix on) have already been effectively ported or adapted to RNA-Seq normalized counts, most of the low-level preprocessing, quality control, and normalization techniques are specific to microarrays, if not even platform-specific (Affymetrix, Agilent, or Illumina). In such a perspective, the risk of an imminent and general loss of the technical know-how needed to analyze expression data from microarray experiments is real.

Nevertheless, we believe that there are a number of good reasons to keep this expertise alive or, even better, to *freeze and package* our current gold standards of microarray data analysis in order to provide present and future researchers with suitable tools to deal with this kind of data at every occurrence:

1. Despite the transcriptomics trends, thousands of articles referring to microarray experiments or microarray analyses are still published yearly [16], and this is likely to continue for a while;
2. The aforementioned GEO and AE databases contain a huge amount of microarray raw data (more than 10^4^ studies overall) that still deserve to be explored by reanalyses (e.g., [17]) or meta-analysis (see e.g., [18]);
3. There is a growing concern about reproducibility of research results in biomedical sciences and bioinformatics [19–22];
4. Typical pipelines for microarray analysis are custom scripts made up of multiple files and several R functions from different Bioconductor packages; dealing with this code – and correctly running a new analysis – months or years later can be source of frustration for many researchers, even among bioinformaticians.

To meet all these needs and make microarray data analysis more accessible to the broad community of biologists, we developed bioTEA (TEA standing for Transcript Expression Analysis), a stand-alone dockerized pipeline for the retrieval, preprocessing, differential expression analysis, and annotation of transcriptomic data from one-color (high-density) microarrays. Its containerized design ensures reproducibility of results over time and across multiple informatics platforms, providing a friendly Command Line Interface (CLI) that frees the end user from coding, and R syntax in particular. Notably, even though specifically designed with microarray technology in mind, bioTEA can handle RNA-Seq data provided in the form of read counts, thus extending its scope of applicability and making it a powerful and invaluable bioinformatics tool.

## 2. Materials and Methods

BioTEA can be used to perform all steps of a classical Differential Expression Analysis (DEA): data retrieval (from GEO), data preprocessing, differential expression analysis by either empirical Bayes moderated *t*-test (i.e., `limma` R package [23]) and/or Rank Product statistics [24,25], and result annotation with metadata. The user is presented with an intuitive, easy-to-use CLI to perform every step of the analysis, which is installed as `biotea`. A standard bioTEA analysis involves the following 5 steps.

1. With `biotea retrieve`, data is directly downloaded from GEO, notably with sample metadata included that will be useful, although not required, to perform successive analysis steps.
2. With `biotea prepare`, raw expression data (as returned by the microarray scanner setup, or as downloaded according to the previous step) is read, parsed, normalized, and quality-controlled. The normalized expression matrix is saved to file, along with Quality-Control (QC) plots.
3. With `biotea initialize`, the differential expression analysis is initialized, optionally by parsing metadata about the samples. This allows the user to quickly set a variety of analysis options such as variable of interest (e.g., treatment status), contrast of interest (e.g., “treated” vs “control”), and analysis batches for batch effect correction. This steps generate an options file that records the various choices made. This allows the user to inspect, edit, or share them for reproducibility.
4. With `biotea analyze`, the option file (such as from the previous step) and the expression matrix is read and DEA is performed. This generates several QC plots, along with differential expression tables as a final result. Conservative filtering is also performed on the data before the analysis, to increase statistical power.
5. With `biotea annotations`, annotations (such as gene symbol or gene name) can be added to expression matrix files or differential expression tables to allow further analyses (such as Gene Ontology (GO) enrichment analysis) and considerations.

It is important to note that these steps need not be run in order, as they are independent of each other. For example, if expression data is already available from other sources, one may directly run `biotea analyze`.

Reproducibility was a key goal in the design of bioTEA. Data preparation, analysis and annotation is performed by code hosted inside a Docker container. This allows for computational reproducibility of these steps, while preserving the automated nature of the tool. Options for the core DEA steps are inputted through an options file, allowing investigators to share the analysis quickly and easily. Indeed, given the same input files and options file, the container assures identical analysis output, independent of operating system version or hardware. Finally, all bioTEA commands log their output in log files, allowing further inspection of run-time events.

### 2.1. The bioTEA container

BioTEA is divided in two intimately related components: the bioTEA docker container and the bioTEA command-line interface. The bioTEA container is structured in modules that perform different functions, all written in the R programming language. A single shared entrypoint binds all modules together. The container performs all the core analysis steps, namely data preparation, DEA, and annotation. The general structure of the container and its modules are shown in Figure 1.

**Figure 1.**
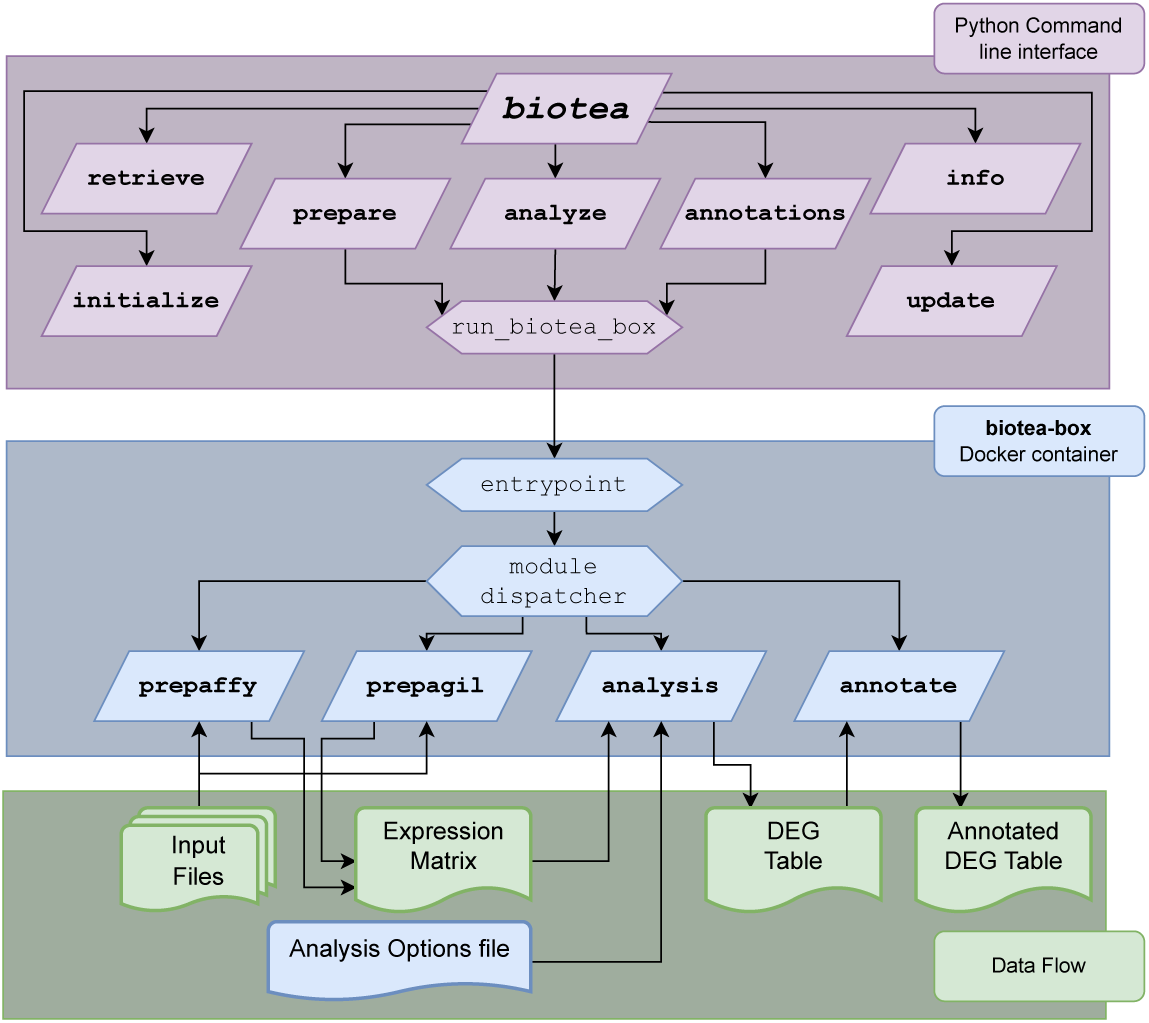
General structure of bioTEA. The python CLI is accessed with the `biotea` command. Launching of the Docker container is handled by a single function, `run_biotea_box`. A shared entrypoint is run upon starting the Docker container, which parses the options and runs the invoked module. File input and output are handled through Docker mounts to predetermined locations inside the running image. In the lower part of the figure, a simplified schema of the input and outputs used by the various modules.

Calling the container directly is complex, as several arguments are needed both to configure the Docker daemon (e.g., input-output mounting path) and the modules themselves. To make this task more accessible to the end user, the bioTEA python package was created. This package exposes the `biotea` command, and handles argument parsing and the correct invocation of the docker daemon. It also provides additional helper commands that are useful in the overall analysis. To promote reproducibility, no data analysis is performed by the bioTEA python package directly.

In the following sections, we aim to explain in more detail the bioTEA container modules, the key pipeline choices we made while writing the core analysis code, and the structure and usage of the bioTEA python package.

### 2.2. Microarray data preparation

The two preparatory modules `prepaffy` and `prepagil` can be used to handle the preprocessing of Affymetrix and Agilent microarray raw data, respectively. As input, both modules take a series of text files generated by the specific microarray scanning apparatus. Then the following steps are performed.

1. Find and load all input files. `prepaffy` searches the target folder for all files ending with the `.CEL` file extension, while `prepagil` searches by default all `.txt` files. However, as `.txt` is a very common file extension, `prepagil` search criteria can be customized in the call. Also, regular expressions are supported by both the modules.
2. Merge all inputs into a single expression data object. Both commands do this automatically using `oligo` and `limma` packages for Affymetrix and Agilent data, respectively.
3. Generate QC plots before normalization. For each sample a Bland-Altman plot (MA plot) is saved along with an overall expression boxplot.
4. Normalize the expression data. Both data types are background-subtracted (through the *normexp* algorithm) and interarray-normalized (through the *quantile-quantile* procedure). Additionally, for Affymetrix data, the Robust Multichip Average (RMA) procedure as provided by the `oligo` package is used to collapse individual probes to the probe-set they belong.
5. Generate new QC plots after normalization. New MA plots and an overall expression boxplot are saved to appreciate the effects of normalization.
6. If specified, remove control probes from the dataset. In particular, negative control probes (i.e., sequences designed to remain unhybridized) represent a sensible estimate for the expected intensity of unexpressed genes as a result of background and nonspecific hybridization signals. Such an estimate is outputted and logged when handling Agilent data, as it can be used as a filter threshold value in the downstream analysis. This step is carried out by bioTEA for Agilent data, and by the `oligo` package for Affymetrix data.
7. Collapse the replicate probes in Agilent arrays. Data is collapsed by taking the mean value of replicate probes found inside each sample.
8. Save the final expression matrix as the output file. The expression matrix is saved in `.csv` format, ready to be analyzed by the other modules of bioTEA, or with customized pipelines.

For more information on the input-output data structures and formats, please refer to Appendix A.

### 2.3. Performing differential expression analysis

The `analysis` module handles the core DEA, which consists in detecting those genes showing different levels of expression across two or more experimental groups, each made up of many biological replicate samples. To this end, samples are labeled using the levels of an input variable named “*experimental design*” (e.g., “treatment1”, “treatment2”, “control”). Differentially Expressed Genes (DEGs) are searched between these groups of samples as indicated by the *contrasts* variable (e.g., “treatment1” vs “control”). Any number of contrasts can be tested in a single run. Beyond a number of parameters—including the experimental design and the contrasts of interest—the module takes as input an expression matrix such as the one generated by the `prep-` modules.

The analysis can be divided into 5 steps: optional (re-)normalization, input visualization, filtering, statistical analysis, output visualization. First, data can be *quantile-quantile* normalized (which is useful only if they were not normalized beforehand) and diagnostic plots are saved before and after this step.

Samples are then clustered by means of a hierarchical algorithm and through Principal Component Analysis (PCA). A dendrogram is saved along with the distribution of the samples in the space of the first principal components. Both representations are useful to spot possible batch effects, which may have a profound effect on the final DEG lists if not properly accounted for. In addition, a mean-variance plot for each experimental condition is included for further QC inspections.

Expression data is then filtered to remove low expression genes and increase the statistical power of the subsequent tests. Filtering is accomplished in a conservative way, according to three parameters: *κ* being the minimum group-wise presence, *θ*_*E*_ being a threshold value on log_2_ expression levels, and *θ*_*F*_ being a threshold on log_2_ fold change values. An entry (a single probe, or probe-set, or transcript cluster, depending on the input type) is retained only if its expression is above *θ*_*E*_ in at least *κ* percentage of samples of at least one experimental group. In addition, while not removed outright, genes that result in |log_2_ *FC*| < *θ*_*F*_ will be marked as “non-differentially expressed” in the final output, even if they show statistical significance, because they usually have little biological importance and can be hard to validate through other means (such as quantitative PCR). However, they are still considered in the computation.

The detection of DEGs is handled by `limma` [23], one of the *de facto* standards for this purpose, and by `RankProd` [24,25], a non-parametric alternative. `Limma` detects DEGs by fitting linear models and performing a gene-wise moderated *t*-test taking advantage of empirical Bayes methods to return more reliable *p*-values even in the case of small sample size. Notably, since `limma` linear models that can take into account a variety of different variables, the user may define a “batch” variable to specify which batch the samples originate from. Batch effect correction is handled during the statistical analysis, both in the `RankProd` and `limma` tests. We chose not to correct the batch effect *a priory*, as recent literature suggested that this may lead to data distortion and wrong conclusions. (PMID: NigaardMethods)

After running `limma` and/or `rankprod`, bioTEA generates a series of diagnostic plots, including MA plots and volcano plots. If both the `limma` and `RankProd` analyses are run, a comparison of the two outputs in the form of Venn diagrams is also included.

The output of the `analysis` module consists in a series of `.csv` format files, one per input contrast per analysis tool (either `RankProd` or `limma` or both), named “DEG tables”. Each table contains every retained entry from the input file, along with information about test statistics, *p*-value, False Discovery Rate (FDR) *q*-value, as well as average expression and log_2_ FC. Also, an additional “markings” column is included labeling each entry as significantly upregulated (1), downregulated (− 1), or non differentially expressed (0). An entry is labeled as differentially expressed if it features a *q*-value strictly less than the critical FDR value of 0.05, and if its average log_2_ FC is greater than *θ*_*F*_ or less than *− θ*_*F*_, as noted before.

If specified in the input, the DEG tables can be automatically annotated and gene symbols are also used to enrich some output plots, such as volcano plots.

As an option, bioTEA can also analyze RNA-seq data in the form of counts, such as those generated from the use of HTSeq [26]. The discrete count data is transformed to continuous expression data using the `voom` function [27] provided by the `limma` package.

### 2.4. Annotating results

The `annotate` module adds additional metadata to either expression matrices or DEG tables. The metadata can be sourced by an internal, static annotation file generated with 39 Human microarray annotation packages present on Bioconductor. This was done for reproducibility reasons, so that the same annotations are applied while using the same bioTEA version. However, packaging all annotations present in Bioconductor, spanning multiple species and microarrays, would render the container too heavy memory-wise. In any case, bioTEA may be configured to download annotation data from Bioconductor at run-time, filling this gap.

Added annotations include gene name and Human Genome Organization (HUGO) gene symbol, as well as which package was sourced to obtain the data, and its version.

### 2.5. The bioTEA command line interface

The bioTEA CLI is an application written in Python 3 and compatible with all UNIX-like operating systems. It allows for easy access to the bioTEA container and for additional features, such as the automatic retrieval of data from GEO. As it is packaged and published on PyPA, installation is straightforward with the standard Python utility `pip`.

The bioTEA CLI handles the download, update, and removal of the bioTEA containers automatically or on-demand of the user.

### 2.6. Source code availability and installation

The bioTEA command line interface is available on PyPI [28], and its code, as well as the code used in the bioTEA container, is available on GitHub [29]. As already noted, a pre-compiled version of the bioTEA image is available at Docker Hub [30].

For reproducibility, bioTEA follows Semantic Versioning 2 for its versioning schema [31]. At the time of writing, the latest bioTEA CLI version is `v1.0.0`, and the current bioTEA image version is `v1.0.1`. The bioTEA versions might change as patches and bug-fixes are introduced in the future. It is important to note that the versions of the two components might diverge; however, we strive to keep compatibility between the two components with identical “major” versions.

To install the tool, install Python version 3.10 or later, as well as the Docker engine and daemon first. Afterwards, install bioTEA with `pip install bioTEA`. The `pip` utility handles the download, installation and update of the tool. Afterwards, the bioTEA CLI may be invoked with the `biotea` command.

## 3. Results

In this section, we exemplified the usage of bioTEA by analyzing a publicly available dataset. This example analysis was run with bioTEA version `0.1.2`. The Docker engine was pre-installed according to the official documentation (server version `20.10.14`). BioTEA was installed with the `pip install biotea` command inside a Python `3.10` virtual environment (bioTEA is compatible with Python version `3.9` and up). The latest bioTEA container version was installed with the `biotea update` command.

### 3.1. Data retrieval

We chose to re-analyze the data provided by Zhang *et al*. [32,33], regarding samples of pancreatic cancer matched with nearby healthy tissues. Data was retrieved directly from GEO (accession GSE28735) with `biotea retrieve`. This command downloads and generates a number of files in the target directory: a `metadata.csv` file, and two sub-directories, a `raw_data` directory with the compressed samples, and a `unpacked_samples` folder with the uncompressed samples.

### 3.2. Preprocessing and quality control

To preprocess the data, uncompressed samples were converted to flat `.CEL` files with `gunzip`. Then, we processed the raw samples to get the expression matrix with `biotea prepare affymetrix`. The result is a series of QC plots, saved in a new `PrepAffy Figures` folder, and an expression matrix file, saved in `.csv` format.

The QC plots are comprised of two MA plots per sample, one before normalization and one after normalization. In each case, the *M* statistic is calculated between the sample and the median value for the same entry in all other samples. The so generated plots are enumerated, starting from the sample that produced the less linear `Loess` fit allowing the user to detect at a glance the samples with lower quality, especially in experiments with very high sample size. Two of these plots can be seen in Figure 2. Additionally, two global boxplots and density plots are generated, before and after data normalization, to appreciate the effects of the normalization steps.

**Figure 2.**
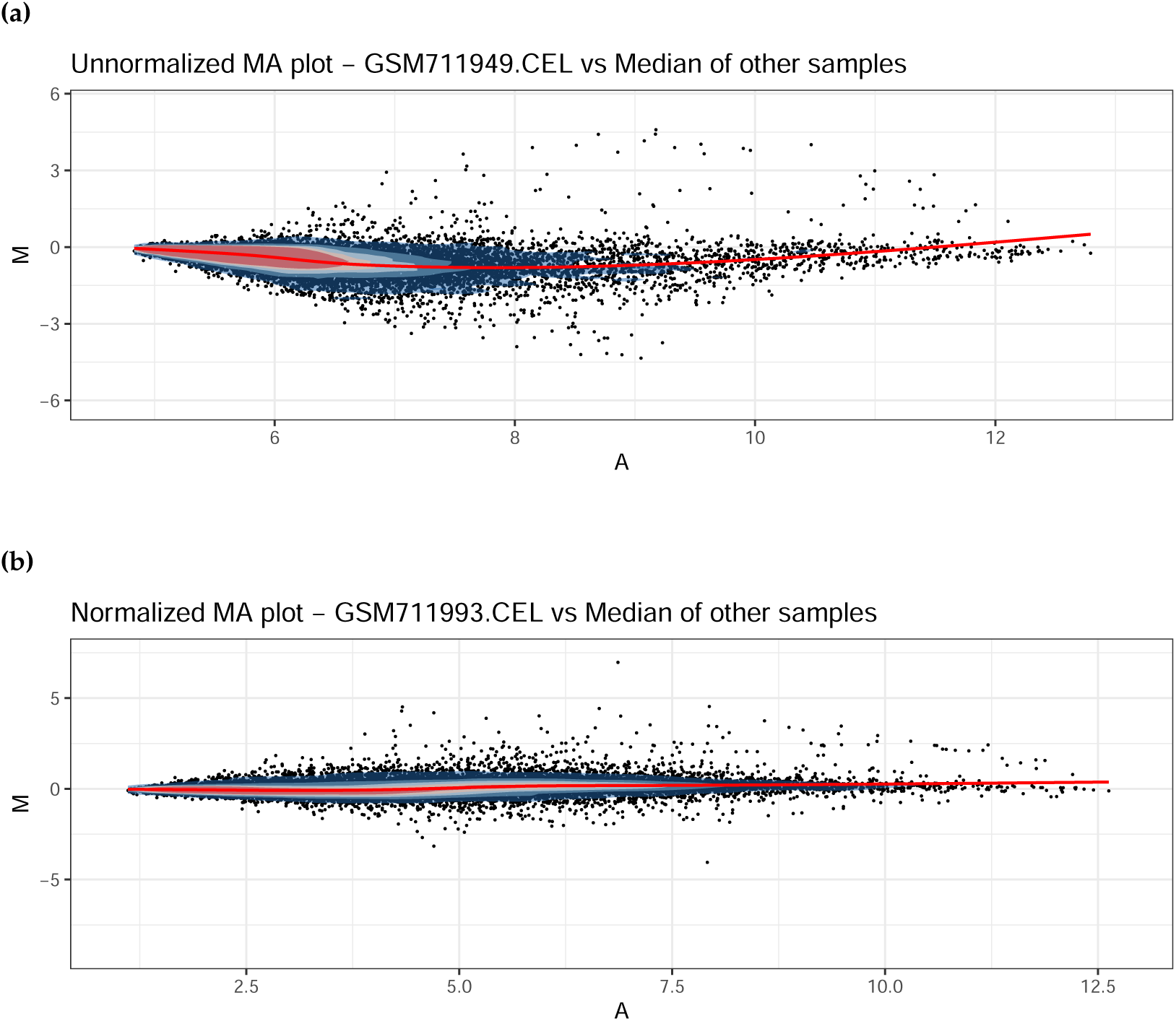
Two Bland-Altman plots showing the distributions of the genes before (a) and after (b) normalization procedures. Data taken from the Zhang dataset.

### 3.3. Differential expression analysis

The option file for `biotea analyze` was generated with `biotea initialize` and then manually edited to set the right parameters for the analysis. The analysis was then launched with `biotea analyze`. The result is a collection of files generated in the target output folder, which includes a series of QC plots, as well as the output DEG tables.

Samples are automatically renamed according to the levels of the main variable of interest, which, in this example, are either “tumor” or “normal”. Importantly, the order of the input samples matters, as the labels are applied to the samples order-wise. For this reason, a correspondence matrix showing the new labels along with the original sample names is saved in the output directory for manual check.

QC plots include a boxplot and a density plot of the input data, similar to those produced by `biotea prepare`, to check that all the samples are properly inter-array normalized. A clustering dendrogram and a PCA plot are also included, to detect possible batch effects (Figure 3). MA plots comparing samples of the different groups of interest are saved. The same plots are additionally saved with DEGs—as detected by both `limma` and RankProduct—overlaid, as shown in Figure 4.

**Figure 3.**
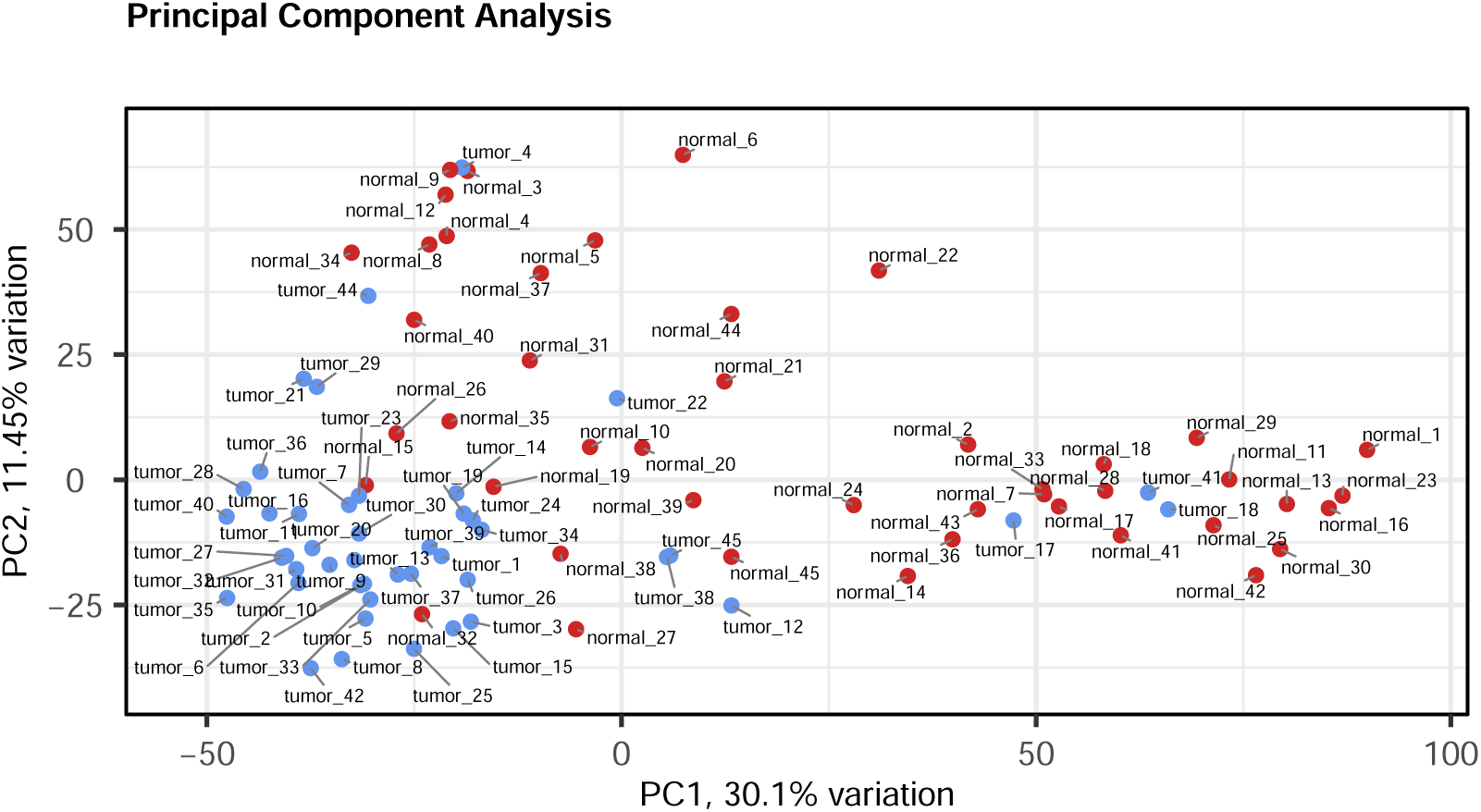
A PCA plot of each samples in the Zhang dataset. The samples are well clustered according to their different sample status (tumor or normal) and no other effect seems to be present. This is a sign of high data quality.

**Figure 4.**
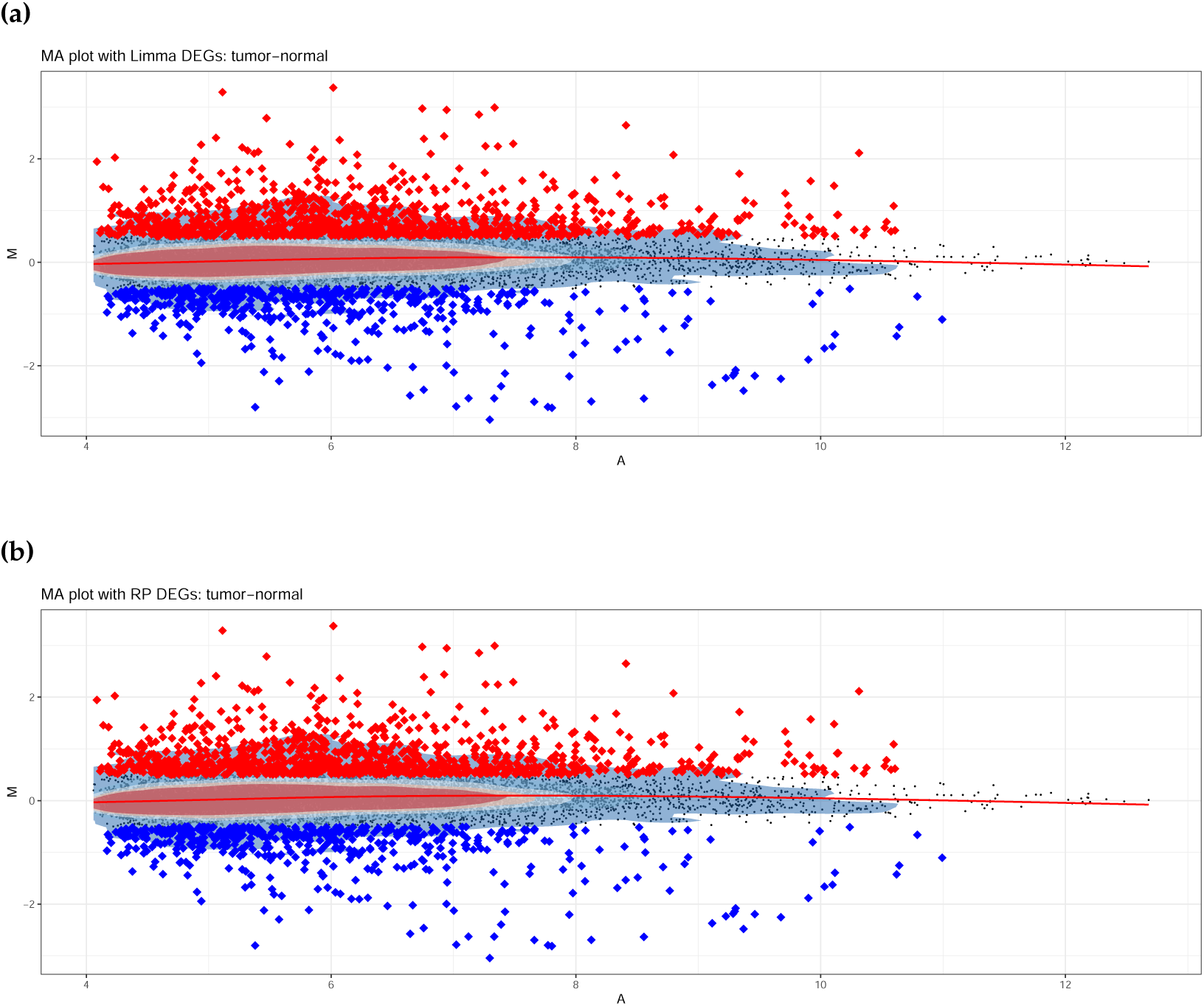
Bland-Altman plots showing the detected DEGs as either upregulated (red) or downregulated (blue), as detected by `limma` (a) or RankProduct (b). The two plots look identical, a sign that DEGs detection was robust.

A very common representation of the results of a DEA is the volcano plot. BioTEA generates one volcano plot per tool per contrasts (see Figure 5).

**Figure 5.**
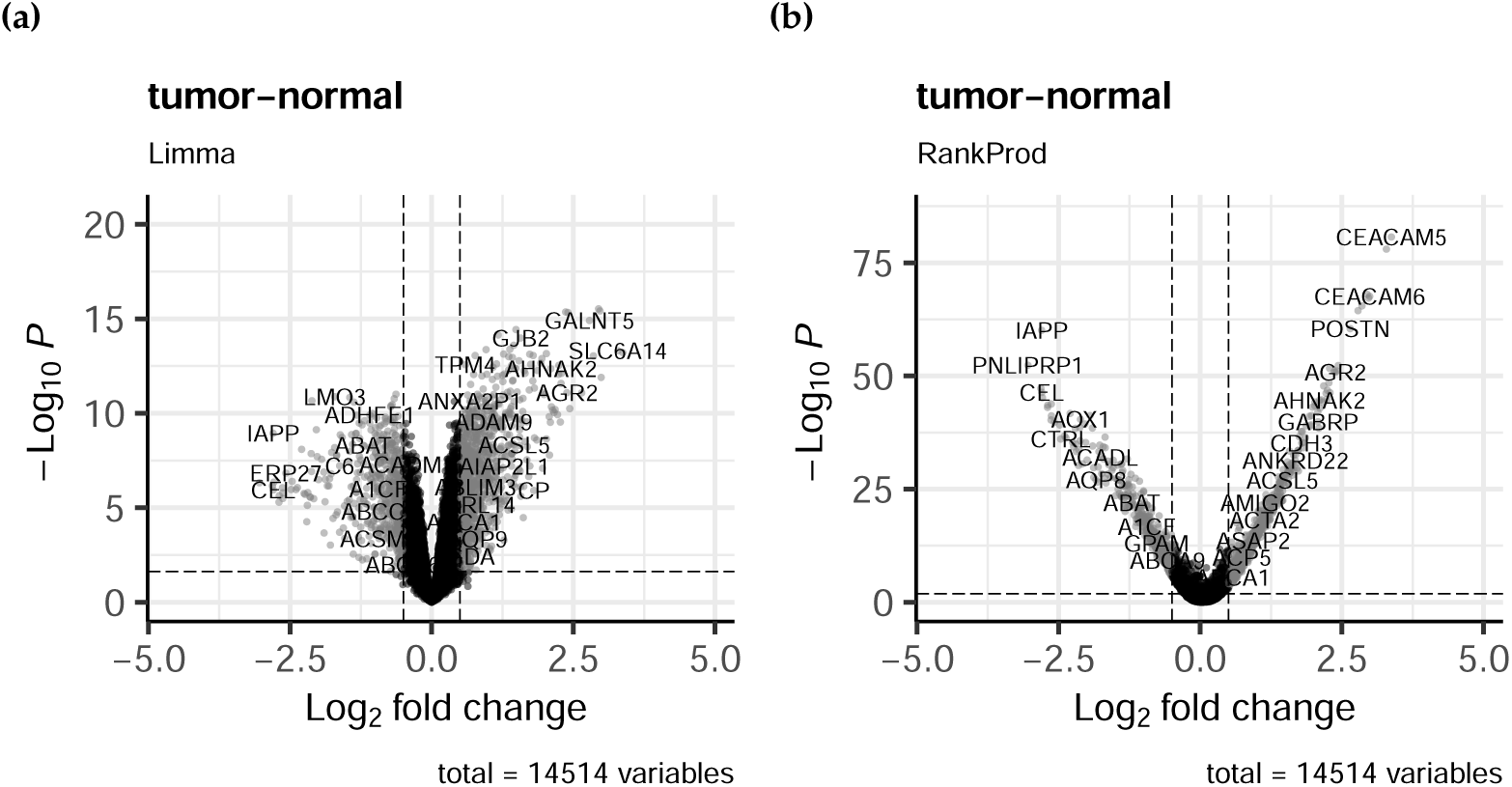
Volcano plots of the DEGs detected by `limma` (a) and RankProduct (b). The different shapes of the distributions are related to how the two algorithms compute the *p*-values. The horizontal dashed line is the *−* log_10_(*p*-value) threshold corresponding to a FDR alpha level of 0.05. The vertical lines are the log_2_ fold-change thresholds, which can be changed by the user and are by default set to 0.5.

The large sample size of the dataset (*n* = 45 for each condition) allowed for high statistical power, reflected in the high number of detected DEGs. Both `limma` and RankProduct detect roughly the same genes as differentially expressed. The detected DEGs were concordant between `limma` and RankProd: both detected the same 70 downregulated genes, while RankProd detected 3 more upregulated DEGs (271) compared to `limma` (268). The shared upregulated genes were identical between the two tools.

## 4. Discussion

In this paper we presented bioTEA as a new tool for the analysis of transcriptomics data. Thanks to the virtualization technology offered by Docker, our software aim at being an OS-independent and robust solution to get reproducible results in differential gene expression analysis. Because of its Python wrapper, bioTEA can be easy installed and run as a high-level command-line application, not requiring any specific programming skill by the user. The main idea behind this project was to use containerization to “hibernate” the most advanced and consolidated standards developed in the last decades for the analysis of microarray data. We think this will allow researchers to preserve the practical ability of mining the huge amount of publicly available microarray datasets, even in the realistic perspective of the forthcoming decline of this technology.

Notably, throughout the software development process, bioTEA has been enriched with many capabilities and features not initially foreseen, including the possibility of analyzing RNA-Seq data, thus greatly extending its scope of applicability. Importantly, although designed to be easy-to-use, bioTEA does not sacrifice analysis power and versatility, the current version of bioTEA providing the following notable features:

- functional reproducibility ensured by Docker container technology
- R/Bioconductor-independent CLI through Python wrapper
- automatic dataset retrieval (from GEO)
- microarray raw data processing capability
- microarray multi-platform support (Agilent and Affymetrix)
- RNA-Seq data support
- automatic annotation for Human arrays (no chip ID needed)
- low-expression gene filtering
- easy experimental design definition by syntax parsing
- two-independent class or paired design testing
- double approach to DEA (parametric by `limma` and non-parametric by RankProduct)
- batch-effect handing
- additional explanatory variable support
- detailed logging
- high quality graphical output
- detailed wiki user-guide
- modular structure suitable for further development

Even if many user-friendly tools for gene expression data analysis have been proposed during the years, either as online services (e.g., GEO2R [34], BART [35]) or locally executable applications (e.g., MeV [36]), in our knowledge none of them encompasses all the bioTEA’s features listed above.

## 5. Conclusions

We think that BioTEA provides an invaluable tool to inexperienced and veteran bioinformaticians alike. For the inexperienced, the ease of use of the tool and its extensive documentation make data preparation and DEAs easy and reproducible. More experienced users will undoubtedly appreciate the simple nature of the output, and the possibility to run many analyses in a short period of time.

To reproduce a bioTEA analysis, it is sufficient to share the commands used and the analysis options file. This renders the analysis process more transparent for the whole community. Additionally, the reproduction of the analysis can be performed even without extensive knowledge of any programming language.

The code for bioTEA is available online and licensed under the permissive MIT license. Users are invited to inspect the code, propose changes to it, or open issues regarding it. The extensive documentation in the form of function doc-strings and comments throughout the code, as well as the modular nature of the code, allow experienced programmers to easily read and modify the codebase.

## Abbreviations

The following abbreviations are used in this manuscript:

AE: ArrayExpress
CLI: Command Line Interface
DEA: Differential Expression Analysis
DEG: Differentially Expressed Gene
FDR: False Discovery Rate
GEO: Gene Expression Omnibus
GO: Gene Ontology
HUGO: Human Genome Organization
MA plot: Bland-Altman plot
NGS: Next Generation Sequencing
PCA: Principal Component Analysis
PFP: Proportion of False Positives
QC: Quality-Control
RMA: Robust Multichip Average.

## Appendix A. Input and output file formats

This section details the input and output file formats used and generated by the tool.

- Expression matrix: Matrix in `.csv` format, with a header row, a column with entry IDs (arbitrarily named `probe_id`, but not necessarily representing actual probe IDs.) and additional columns, one per input file, with the respective expression data (with arbitrary names). Each row in the matrix represents the expression of a single entry.
- DEG table: Matrix in `.csv` format, with a series of columns:

– The `probe_id` column contains the entry ids of the data (probe IDs or ENSEMBL IDs);
– The `LogFC` column lists the Fold Change value of that entry, in the contrast of interest.
– The `AveExprs` column contains the arithmetic mean of the expression values of the entry between all samples.
– The `t` column reports the t-statistic computed by `limma`.
– The `P.Value` column holds the raw P-values computed by `limma`.
– The `adj.P.Val` column holds the P-values computed by `limma` and adjusted with the Benjamini–Hochberg procedure, producing FDR values.
– The `B` column contains the Bayes factor for the entry. Only computed by `limma`.
– The `gene.index` column contains gene idexes as computed by RankProduct. Only present in the DEG table computed by RankProduct.
– The `RP/Rsum.UP` and `RP/Rsum.DOWN` contain the RankProduct statistics for each entry. Only present in the DEG table computed by RankProduct.
– `pfp.UP` and `pfp.DOWN` are only present in the RankProduct output. The Proportion of False Positives (PFP) of up- and down-regulated genes, respectively.
– `P.Value.UP` and `P.Value.DOWN` are only present in the RankProduct output. The P-values for up- and down-regulated genes, respectively.
– The `markings` column marks as upregulated (1), downregulated (− 1) or non-DEG (0) each entry. For `limma`, this is done by the `decidetests` function, provided by `limma`, with a P-value threshold of 0.05. Additionally, genes with a lower Fold Changes than what specified by the user are marked as non differentially expressed. For RankProduct, the markings are similarly applied. Genes that show a PFP value of less than 0.05 are marked as up- or down-regulated (looking at the `pfp.UP` and `pfp.DOWN` values, respectively), and genes that show a fold change lower than what specified are marked as non differentially expressed. Therefore, this column can be used to quickly subset the output.

Annotation can be applied on any Expression Matrix or DEG table, as the only requirement is the presence of a `probe_id` column. The added metadata columns are always three: `SYMBOL` (with the HUGO gene symbol), `GENENAME` (the long gene name) and `ENSEMBL` (with the Ensemble id related to the gene). Additionally, a `package_name` and `version` column list the package(s) used to source the annotations and its version(s) are added.

## References

1. Duggan, D.J.; Bittner, M.; Chen, Y.; Meltzer, P.; Trent, J.M. Expression Profiling Using cDNA Microarrays. Nature Genetics 1999, 21, 10–14. doi:10.1038/4434.

2. Schena, M.; Shalon, D.; Davis, R.W.; Brown, P.O. Quantitative Monitoring of Gene Expression Patterns with a Complementary DNA Microarray. Science (New York, N.Y.) 1995, 270, 467–470. doi:10.1126/science.270.5235.467.

3. Shalon, D.; Smith, S.J.; Brown, P.O. A DNA Microarray System for Analyzing Complex DNA Samples Using Two-Color Fluorescent Probe Hybridization. Genome Research 1996, 6, 639–645. doi:10.1101/gr.6.7.639.

4. Larkin, J.E.; Frank, B.C.; Gavras, H.; Sultana, R.; Quackenbush, J. Independence and Reproducibility across Microarray Platforms. Nature Methods 2005, 2, 337–344. doi:10.1038/nmeth757.

5. Tarca, A.L.; Romero, R.; Draghici, S. Analysis of Microarray Experiments of Gene Expression Profiling. American Journal of Obstetrics and Gynecology 2006, 195, 373–388. doi:10.1016/j.ajog.2006.07.001.

6. Mayo, M.S.; Gajewski, B.J.; Morris, J.S. Some Statistical Issues in Microarray Gene Expression Data. Radiation Research 2006, 165, 745–748. doi:10.1667/RR3576.1.

7. Verducci, J.S.; Melfi, V.F.; Lin, S.; Wang, Z.; Roy, S.; Sen, C.K. Microarray Analysis of Gene Expression: Considerations in Data Mining and Statistical Treatment. Physiological Genomics 2006, 25, 355–363. doi:10.1152/physiolgenomics.00314.2004.

8. Slonim, D.K.; Yanai, I. Getting Started in Gene Expression Microarray Analysis. PLoS computational biology 2009, 5, e1000543. doi:10.1371/journal.pcbi.1000543.

9. Chen, J.J. Key Aspects of Analyzing Microarray Gene-Expression Data. Pharmacogenomics 2007, 8, 473–482. doi:10.2217/14622416.8.5.473.

10. Gentleman, R.C.; Carey, V.J.; Bates, D.M.; Bolstad, B.; Dettling, M.; Dudoit, S.; Ellis, B.; Gautier, L.; Ge, Y.; Gentry, J.; et al. Bioconductor: Open Software Development for Computational Biology and Bioinformatics. Genome Biology 2004, 5, R80. doi:10.1186/gb-2004-5-10-r80.

11. Home - GEO - NCBI. https://www.ncbi.nlm.nih.gov/geo/.

12. Edgar, R.; Domrachev, M.; Lash, A.E. Gene Expression Omnibus: NCBI Gene Expression and Hybridization Array Data Repository. Nucleic Acids Research 2002, 30, 207–210. doi:10.1093/nar/30.1.207.

13. Browse < ArrayExpress < EMBL-EBI. https://www.ebi.ac.uk/arrayexpress/browse.html.

14. Brazma, A.; Parkinson, H.; Sarkans, U.; Shojatalab, M.; Vilo, J.; Abeygunawardena, N.; Holloway, E.; Kapushesky, M.; Kemmeren, P.; Lara, G.G.; et al. ArrayExpress–a Public Repository for Microarray Gene Expression Data at the EBI. Nucleic Acids Research 2003, 31, 68–71. doi:10.1093/nar/gkg091.

15. Wang, Z.; Gerstein, M.; Snyder, M. RNA-Seq: A Revolutionary Tool for Transcriptomics. Nature Reviews. Genetics 2009, 10, 57–63. doi:10.1038/nrg2484.

16. Lowe, R.; Shirley, N.; Bleackley, M.; Dolan, S.; Shafee, T. Transcriptomics Technologies. PLoS computational biology 2017, 13, e1005457. doi:10.1371/journal.pcbi.1005457.

17. Sabaie, H.; Mazaheri Moghaddam, M.; Mazaheri Moghaddam, M.; Amirinejad, N.; Asadi, M.R.; Daneshmandpour, Y.; Hussen, B.M.; Taheri, M.; Rezazadeh, M. Long Non-Coding RNA-associated Competing Endogenous RNA Axes in the Olfactory Epithelium in Schizophrenia: A Bioinformatics Analysis. Scientific Reports 2021, 11, 24497. doi:10.1038/s41598-021-04326-0.

18. Leal-Calvo, T.; Moraes, M.O. Reanalysis and Integration of Public Microarray Datasets Reveals Novel Host Genes Modulated in Leprosy. Molecular genetics and genomics: MGG 2020, 295, 1355–1368. doi:10.1007/s00438-020-01705-6.

19. Baker, M. 1,500 Scientists Lift the Lid on Reproducibility. Nature 2016, 533, 452–454. doi:10.1038/533452a.

20. Begley, C.G.; Ellis, L.M. Drug Development: Raise Standards for Preclinical Cancer Research. Nature 2012, 483, 531–533. doi:10.1038/483531a.

21. Samsa, G.; Samsa, L. A Guide to Reproducibility in Preclinical Research. Academic Medicine: Journal of the Association of American Medical Colleges 2019, 94, 47–52. doi:10.1097/ACM.0000000000002351.

22. Sandve, G.K.; Nekrutenko, A.; Taylor, J.; Hovig, E. Ten Simple Rules for Reproducible Computational Research. PLoS computational biology 2013, 9, e1003285. doi:10.1371/journal.pcbi.1003285.

23. Ritchie, M.E.; Phipson, B.; Wu, D.; Hu, Y.; Law, C.W.; Shi, W.; Smyth, G.K. Limma Powers Differential Expression Analyses for RNA-sequencing and Microarray Studies. Nucleic Acids Research 2015, 43, e47. https://doi.org/10.1093/nar/gkv007, doi:10.1093/nar/gkv007.

24. Hong, F.; Breitling, R.; McEntee, C.W.; Wittner, B.S.; Nemhauser, J.L.; Chory, J. RankProd: A Bioconductor Package for Detecting Differentially Expressed Genes in Meta-Analysis. Bioinformatics (Oxford, England) 2006, 22, 2825–2827. doi:10.1093/bioinformatics/btl476.

25. Del Carratore, F.; Jankevics, A.; Eisinga, R.; Heskes, T.; Hong, F.; Breitling, R. RankProd 2.0: A Refactored Bioconductor Package for Detecting Differentially Expressed Features in Molecular Profiling Datasets. Bioinformatics 2017, 33, 2774–2775. https://www.ncbi.nlm.nih.gov/pmc/articles/PMC5860065/, doi:10.1093/bioinformatics/btx292.

26. Putri, G.H.; Anders, S.; Pyl, P.T.; Pimanda, J.E.; Zanini, F. Analysing High-Throughput Sequencing Data in Python with HTSeq 2.0. Bioinformatics 2022, p. btac166. https://doi.org/10.1093/bioinformatics/btac166, doi:10.1093/bioinformatics/btac166.

27. Law, C.W.; Chen, Y.; Shi, W.; Smyth, G.K. Voom: Precision Weights Unlock Linear Model Analysis Tools for RNA-seq Read Counts. Genome Biology 2014, 15, R29. https://doi.org/10.1186/gb-2014-15-2-r29, doi:10.1186/gb-2014-15-2-r29.

28. bioTEA · PyPI. https://pypi.org/project/bioTEA/.

29. BioTEA. CMA-Lab, 2022. https://github.com/CMA-Lab/bioTEA.

30. Cmalabscience/Biotea-Box Tags | Docker Hub. https://hub.docker.com/r/cmalabscience/biotea-box/tags.

31. Preston-Werner, T. Semantic Versioning 2.0.0. https://semver.org/.

32. Zhang, G.; Schetter, A.; He, P.; Funamizu, N.; Gaedcke, J.; Ghadimi, B.M.; Ried, T.; Hassan, R.; Yfantis, H.G.; Lee, D.H.; et al. DPEP1 Inhibits Tumor Cell Invasiveness, Enhances Chemosensitivity and Predicts Clinical Outcome in Pancreatic Ductal Adenocarcinoma. PloS One 2012, 7, e31507. doi:10.1371/journal.pone.0031507.

33. Zhang, G.; He, P.; Tan, H.; Budhu, A.; Gaedcke, J.; Ghadimi, B.M.; Ried, T.; Yfantis, H.G.; Lee, D.H.; Maitra, A.; et al. Integration of Metabolomics and Transcriptomics Revealed a Fatty Acid Network Exerting Growth Inhibitory Effects in Human Pancreatic Cancer. Clinical Cancer Research: An Official Journal of the American Association for Cancer Research 2013, 19, 4983–4993. doi:10.1158/1078-0432.CCR-13-0209.

34. Barrett, T.; Wilhite, S.E.; Ledoux, P.; Evangelista, C.; Kim, I.F.; Tomashevsky, M.; Marshall, K.A.; Phillippy, K.H.; Sherman, P.M.; Holko, M.; et al. NCBI GEO: archive for functional genomics data sets–update. Nucleic acids research 2013, 41, D991–5. doi:10.1093/nar/gks1193.

35. Amaral, M.L.; Erikson, G.A.; Shokhirev, M.N. BART: bioinformatics array research tool. BMC Bioinformatics 2018, 19, 296. doi:10.1186/s12859-018-2308-x.

36. Howe, E.A.; Sinha, R.; Schlauch, D.; Quackenbush, J. RNA-Seq analysis in MeV. Bioinformatics 2011, 27, 3209–3210. doi:10.1093/bioinformatics/btr490.

